# Persistent features of intermittent transcription

**DOI:** 10.1101/650895

**Authors:** Michael Wilkinson, Spyros Darmanis, Angela Oliveira Pisco, Greg Huber

**Affiliations:** Chan Zuckerberg Biohub, 499 Illinois Street, San Francisco, CA 94158, USA; School of Mathematics and Statistics, The Open University, Walton Hall, Milton Keynes, MK7 6AA, England

## Abstract

Here we report statistical studies of single-cell mRNA counts from cells derived from different tissues of adult mice. By examining correlations between mRNA gene counts we find strong evidence that when genes are only observed in a small fraction of cells, this is as a consequence of intermittent transcription rather than of expression only in specialized cell types. Count statistics are used to estimate a peak transcription level for each gene, and a probability for the gene to be active in any given cell. We find that the peak transcription levels are approximately constant across different tissue types, but the gene expression probabilities may be markedly different. Both these quantities have very wide ranges of values, with a probability density function well approximated by a power law.

**Author summary:** Using evidence from single-cell mRNA counts, we argue that the expression of many genes in individual mouse cells is highly intermittent. Comparing cells from different tissues, we find that the peak activity of a given gene is approximately the same in all tissue types, whereas the probability of a gene being active can differ markedly.

## Introduction

Using nucleic acid polymerase technologies, it has become possible to quantify mRNA transcription processes with ever greater sensitivity [1]. Recently, it has become possible to obtain quantitatively reliable counts of individual gene transcriptions from single cells [2]. This technology has the potential to reveal new insights into the mechanisms and organization of transcription processes. This paper reports on a statistical analysis of the *Tabula Muris* dataset of mRNA transcriptions from individual mouse cells, derived from a range of distinct tissue types [3].

The number of ‘reads’ of mRNA from individual cells is highly variable, with many cells yielding a zero count for a particular gene, while a few cells from the same tissue type might yield hundreds or even thousands of reads. This variation is often ascribed to ‘dropouts’, viewed as a technical consequence of the stochastic nature of the DNA polymerization reaction. However, the variability of the counts is markedly different between different genes, and for many genes the counts are much more variable than those of exogenous RNA sequences which are ‘spiked in’ with known concentrations. The variability cannot, therefore, be explained solely as a technical artifact, and we should therefore consider other, biological interpretations.

In this paper we use single-cell mRNA counts to estimate the probability *p* that each gene is being expressed in a given cell. These probabilities vary greatly, and we find that a significant fraction of genes are expressed with very low probability. There are at least two possible explanations for the wide variability of the gene *expression probability p*, illustrated schematically in figure 1:

- **Case A:** the cell population could be inhomogeneous, with cells from a given tissue differentiating into different types, which express the same set of genes continuously. A small value of *p* is a consequence of a gene being expressed in a rare, specialized cell type. In the schematic of figure 1(a) genes with labels *i* and *j* are expressed by different types of rare cells.
- **Case B:** It could be that the activity of a cell within a given tissue-type population is time-dependent. In this case a small value of *p* is interpreted as an indication that the gene is turned off for most of the time. In the schematic illustration figure 1(b), a cell which is actively expressing a gene has a count equal to a *peak activity α*, but for most of the time the gene is not being transcribed.

**Fig 1.**
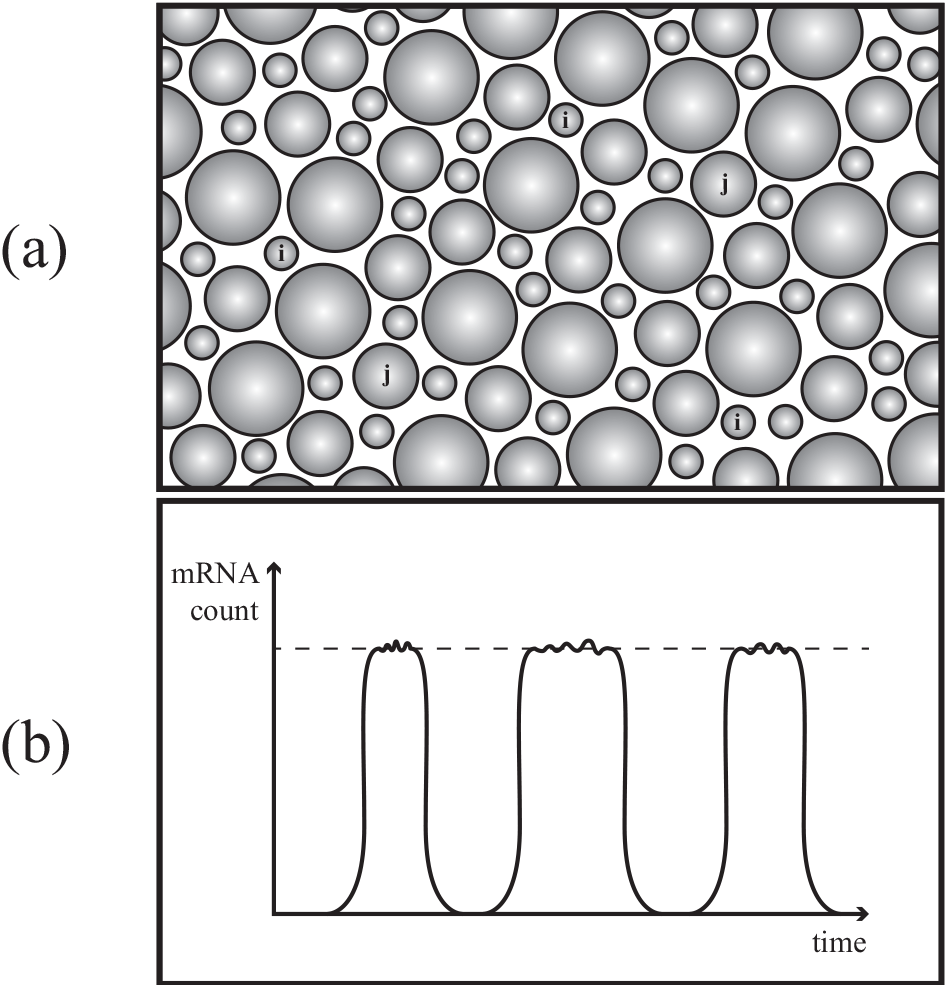
Schematic contrasting two explanations for genes being observed with low probability in single cell mRNA counts. (a) Genes *i* and *j* may be transcribed continuously in rare, specialized cell types. (b) Genes may be intermittently active in all cells.

It is desirable to distinguish clearly between these possibilities. This can be done by considering the coefficients of correlation for expression of different genes in the same cell. The evidence (discussed in section) strongly favors case B above as a model for explaining the occurrence of genes which are expressed with a low probability.

Earlier studies of the time dependence of the expression of single genes have shown evidence that transcription of mRNA occurs in ‘bursts’. This comes from direct observation of the time-dependence of mRNA transcription [4, 5]. It has been remarked that indirect evidence for bursting also comes from from the observation that the variance of single-cell mRNA reads is typically much larger than the mean [5, 6]. The ‘bursty’ transcription is ascribed to being a consequence of a stochastic process, involving binding and un-binding of transcription factors from the DNA [7–9]. More recently, by fitting the statistics of single-cell mRNA reads to stochastic models of bursty transcription [10], burst sizes and burst frequencies have been ascribed to individual genes [11]. The work of Larsson *et al* [11] emphasizes correlations between the kinetic parameters of the bursting model and aspects of the structure of the gene (such as its size) or of its promoters (such as TATA elements). In the concluding discussion of our paper we contrast the use of the bursting model with our own approach.

Because ‘bursting’ transcription has an association with a particular stochastic model which may not be the only viable explanation of the phenomena, we shall refer to *intermittent* transcription in this text. Further information, such as investigation of correlations between gene expression, are required to distinguish between intermittent transcription and gene expression by specialist cells (that is, between cases A and B above). The analysis of the *Tabula Muris* dataset presented here supports the view that intermittent transcription is an ubiquitous phenomenon, extending to genes which are only expressed at very low levels. The fact that mRNA counts are highly variable implies that genes are being turned on and off very slowly, on a timescale much longer than the lifetime of mRNA (which appears to be at least one hour in most mammalian cells [12]).

Because the *Tabula Muris* data set contains information from a range of different tissue types, we are able to assess the effects of cellular differentiation on gene expression. We find evidence that the peak rate of transcription, *α*, is a *persistent* attribute of a gene, which takes similar values in all of the tissues that were surveyed. The probability of transcription is found to be much more variable. These observations are consistent with gene expression being controlled by turning genes on and off, rather than via continuous regulation of their rate of transcription.

Both the peak transcription level *α* and the probability of expression *p* vary over a large range. We also find evidence that the probability density of *α* is well approximated by a power law. The probability density of the gene expression probability varies significantly between different tissues. In the concluding section we discuss whether intermittent transcription is necessarily a stochastic phenomenon, as is frequently supposed [7–9], and why it might confer advantages in organizing activity within a cell.

## A two-parameter characterization of genes

### The Tabula Muris data

The *Tabula Muris* dataset [13] combines single-cell mRNA count data from two different technologies: microfluidic droplet-based 3’-end counting, which provides a survey of thousands of cells per organ at relatively low coverage, and FACS-based full length transcript analysis, which provides higher sensitivity and coverage [3]. The ‘droplet’ data uses unique molecular identifier (UMI) sequences to label individual mRNA molecules, providing a precise assay of the number of mRNA molecules from a given gene that have been amplified. The ‘FACS’ data set records the number of reads after amplification, without counting the number of individual molecules which have been captured. We processed both data set finding that, while not quantitatively equivalent, they are sufficiently consistent to justify our qualitative conclusions.

The FACS-based dataset lists the counts of mRNA molecules obtained from individual cells, for 23,433 protein-coding genes (as annotated in the dataset [13]), obtained from samples of 17 different tissues: the number of sampled cells ranges from 866 (kidney) to 6,007 (heart). In addition to the endogenous mRNA, the samples from each cell were ‘spiked’ with known concentrations of synthetic mRNA, with 96 different combinations of sequences and concentrations. The dataset also contains the number of reads for each of these ‘spike-in’ sequences. The droplet data was obtained from samples of 12 different tissue types: there were a total of 55,638 cells, ranging from 624 heart cells to 11,258 trachea cells.

In the case of the FACS data we only included genes where more than 500 total counts were recorded, neglecting counts of less than 10 in individual cells. In the case of the droplet data, we applied a quality threshold of at least 1000 total reads and at least 500 genes detected.

### Gene parameters

The data lists the number of reads *M*_*ijk*_ of mRNA for a gene with index *i*, in a cell with index *j*, from a tissue with index *k*. The polymerization process is inherently unstable and the experimental parameters may not be exactly the same for all cells, and it was found that the total count for a cell, ∑_*i*_*M*_*ijk*_, varies over a large range (approximately two decades). For this reason, we considered ‘normalizing’ the counts by dividing *M*_*ijk*_ by the total number of counts for each cell (which would be appropriate if our studies were aimed, for example, at characterizing differences between different tissue types). However, evidence from the ‘spike-in’ sequences salted into the FACS data (discussed in detail in section below) indicates that most of the variability of the total count for individual cells is real, rather than a technical artifact. For this reason, we did not normalize the counts.

The count data was processed to produce two statistics for each gene, denoted by *α* and *p*. The *α* variable is a measure of the peak level of transcription of a gene, and the *p* variable characterizes the probability that a given cell will be expressing that gene. If *N*_*k*_ is the number of cells for tissue type *k*, for a given gene with index *i*, we calculate the mean *µ*_*ik*_ and variance 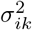 of the counts *M*_*ijk*_, defined by

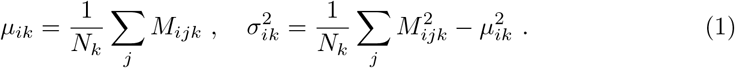

From the means and variances we can construct the quantities

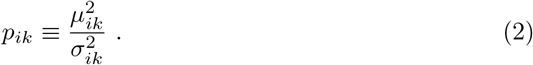

Let us consider what value of *p*_*ik*_ would be expected according to a model where the transcription process is intermittent, in the sense that it is either ‘on’, with a small probability *p*, or else ‘off’, and where the ‘on’ state results in a count equal to *α*. According to this model, the mean and variance would be *µ* = *αp* and *σ*^2^ = *p*(1 − *p*)*α*^2^, so that if *p ≪* 1 then *p ≈ µ*^2^/*σ*^2^, in agreement with equation (2). This indicates that if the transcription occurs intermittently, with the probability of being ‘on’ being *p ≪* 1, then *p*_*ik*_is a measure of the probability of a cell expressing gene *i* in tissue *k* at a significant rate. Note that equation (2) was motivated by a simple model, under the assumption that *p* is small. It is not intended to be accurate when *p* is close to unity. We remark that the reciprocal quantity *σ*^2^/*µ*^2^ has previously been used as an estimator for ‘noise’ in studies of protein transcription [14].

If the number of counts when a gene is active is *α*, then the mean count is equal to *µ* = *αp*. Given an estimate of *p*, equation (2), we can then estimate *α* by writing *α* = *µ/p*. This indicates that the quantity

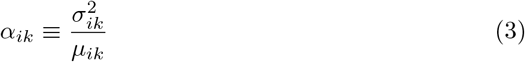

is a measure of the peak level of transcription of gene *i* in tissue type *k*. Thus every gene *i* in tissue *k* can be characterized by two parameters: a probability that it is active, *p*_*ik*_, and a peak activity level *α*_*ik*_, across different tissue types, labeled by *k*. If, when transcription of a gene is ‘turned on’, mRNA is produced at a rate *R*_p_ and destroyed at a rate *R*_d_, then the number of UMIs in the droplet data is expected to be *α ~ R*_p_*/R*_d_. We cannot distinguish directly whether the variability in *α* is primarily due to variations in the rate of production or the rate of destruction.

Both quantities, *p*_*ik*_ and *α*_*ik*_, have a broad range of values spanning two and three decades respectively. For this reason, it is useful to use logarithmic variables

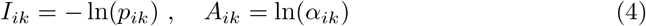

so that the statistics are not dominated by properties of the largest values. The negative sign in the definition of *I* ensures that this variable takes positive values. The *A*_*ik*_ will be referred to as the *activity* of gene *i* in tissue *k*, and *I*_*ik*_ will be termed its *intermittency*.

### Characterization of spiked-in sequences

There were 96 exogenous sequences (denoted by labels ERCC-000*xx*, where *xx* is a two-digit number) ‘spiked’ into the FACS samples with known concentrations. These were used to give an indication of the reproducibility of the experimental procedure. Because the PCR process is unstable, it amplifies fluctuations, such as stochastic variation of counts or small differences in the experimental parameters between reaction cells. For this reason, it may be advantageous to ‘normalize’ the counts, replacing *M*_*ijk*_ by

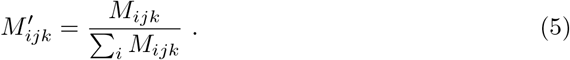

An alternative scenario is that the total number of mRNA molecules in a cell may have large variations, which are not due to measurement errors. In this case the measurements would be distorted by applying the normalization. We used the spike-in sequences to provide a test of whether it is appropriate to normalize the data. Fluctuations of the logarithm of the counts of spike-in sequences give an indication of the fractional errors. We compared the variance of ln *M* (un-normalized) to the variance of ln *M′* (normalized counts) for the exogenous sequences, and found that the variance of the latter was considerably larger (approximately five time higher than the variance of the un-normalized counts). This was mainly due to cells with very low total counts, which cause the genes which are present to be greatly exaggerated if normalization is carried out. Even after eliminating cells in the lower quartile of the total count, the variance of ln(*M′*) was still substantially greater than that of ln(*M*). For this reason, our statistics used the raw (un-normalized) count data.

Because a given exogenous sequence is spiked into all cells at the same concentration, it should, ideally, have zero intermittency. In practice, because of the sources of technical variability mentioned above, the spike-in sequences have non-zero and apparently random values of the intermittency *I*_*ik*_. We find few of the spike-ins which are present at higher concentrations (above 200 amol/*µ*l) give values of the intermittency parameter greater than ln(5) (corresponding to a gene being active with probability less than *p* = 0.2). Accordingly, we regard all genes with *I >* ln(5) as being intermittent.

While other groups have proposed quite complex schemes for normalizing single-cell mRNA counts [15, 16], the use of normalization for the purposes of the present study did not appear to be advantageous.

## Statistical observations and interpretations

### Gene activity correlations

In order to distinguish between two possible models for explaining small values of *p*, cases **A** and **B** discussed in the Introduction and illustrated in figure 1, we used the *Tubula Muris* dataset to examine correlation coefficients of the gene activities: these are

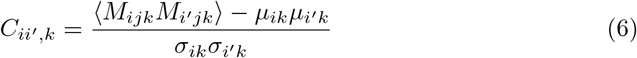

where the angle brackets denote an average over the set of cells in tissue type *k*. Figure 2 shows the probability density function (PDF) of the correlation coefficients for heart and liver tissue, using the droplet data (other tissues give similar results). The distributions of both the positive and the negative correlation coefficients are displayed. Most of the correlation coefficients are extremely small, and not statistically significant. Including all gene pairs, positive and negative coefficients occurred in roughly equal numbers, but figure 2 shows that among the larger correlation coefficients, there is a much smaller proportion of negative coefficients.

**Fig 2.**
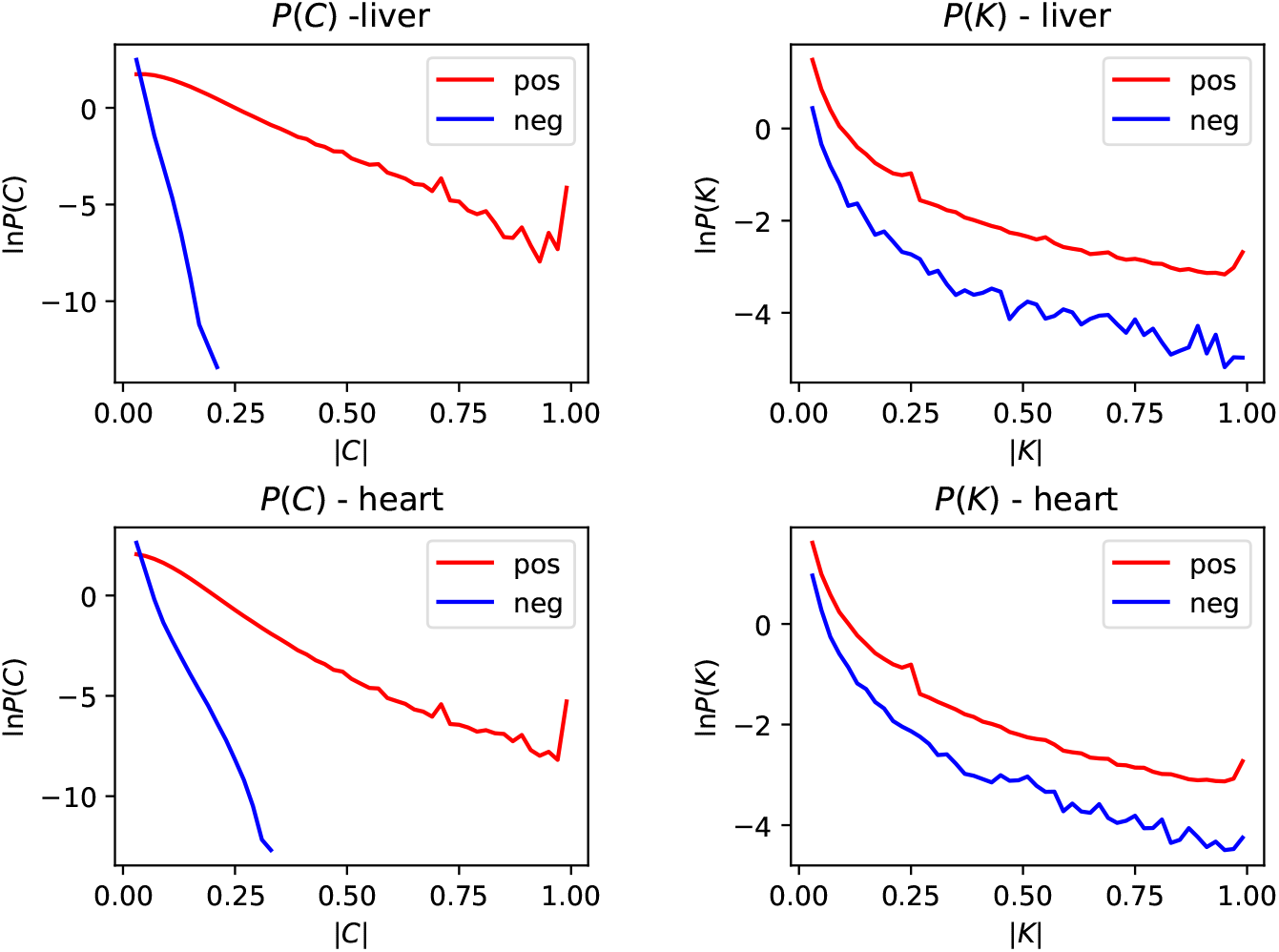
Distribution of the magnitude of gene-activity correlation coefficients, *C* (equation (6)), and of the transformed correlation coefficients, *K* (equation (7)), for both heart and liver cells. In each case we show the distributions for both positive and negative correlations. In cases where there is a significant correlation of gene activity, positive correlations are far more common than negative ones. If genes were active in different cells, the *K* coefficients would be expected to equal −1.

We now argue that these data distinguish between cellular differentiation (case **A**) and intermittent transcription (case **B**). Let us assume that low-expression genes, labelled by index numbers *i* and *i′*, are only expressed in different rare cell types, C_1_ and C_2_ respectively. This implies that if gene *i* is being expressed, so that *M*_*ijk*_ > 0, then the cell is of type C_1_, so that gene *i′* is not expressed, implying that *M_i′ jk_* = 0. Similarly, if *M_i_′ _jk_ >* 0 we are dealing with a cell of type C_2_, and *M*_*ijk*_ = 0. If rarely expressed genes result from cellular differentiation, then pairs of rare genes would not usually be expressed in the same cell, implying that 〈*M_ijk_M_i_′ _jk_*〉 would be equal to zero, because the counts *M*_*ijk*_ would never be non-zero in the same cell for different genes. This implies that the correlation coefficient, given in equation (6) would be negative, and equal to −*µ_ik_µ_i_′_k_/σ_ik_σ_i_′ _k_*. According to the cellular differentiation model, many values of

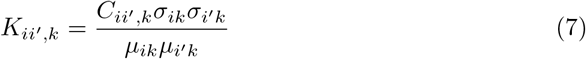

would be very close to −1. In figure 2 we display the PDF of the positive and negative vales of *K_ii′,k_* for heart and liver droplet data. There are very few values of *K_ii′,k_* close to −1. This strongly favors the intermittent transcription model.

### Variation between tissue types

For each gene, we can also define the average of the activity and of the intermittency across different tissues, denoting these by 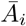 and 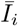 respectively (we used a simple average, rather than one weighted by the numbers of cells in each tissue). It is interesting to consider the fluctuations of activity and intermittency between different tissue types, described by the following quantities:

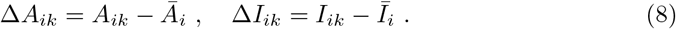

Figure 3 shows ‘heatmaps’ (that is, a 2d histogram color coded for the occupancy of the bins) showing the density of ∆*I*, ∆*A* across all combinations of tissues and genes (these are plotted separately for the two experimental protocols). The most significant feature is that the values of the activity fluctuations ∆*A* show significantly smaller dispersion, than the variation of the intermittency, ∆*I*. The variances of the data points generating figure 3 are Var(∆*A*) = 0.157 and Var(∆*I*) = 1.28 respectively for the droplet data, and Var(∆*A*) = 0.457 and Var(∆*I*) = 1.02 for the FACS data. Another interesting feature is that there are two distinct populations of genes: the bright spot at the center of the heatmap shows that many genes show little variation in their intermittency between different tissues, whereas others show a marked variation in their probability of expression. The surprisingly low value of the variance of ∆*A*_*ik*_ in the droplet data is partly due to there being a significant number of genes with just one count in one cell in this data set.

**Fig 3.**
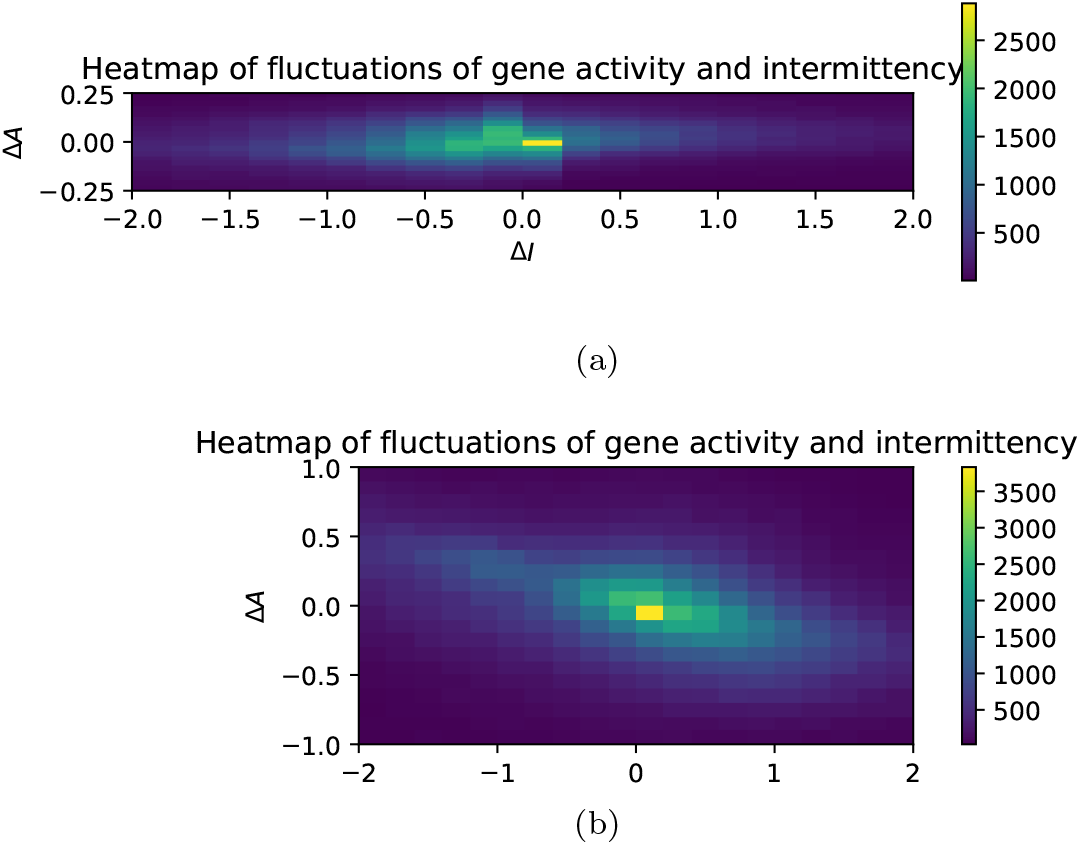
‘Heatmap’ showing the dispersion of the gene activity and intermittency, ∆*A* and ∆*I*, across different tissues. The activity of a gene is almost constant for all tissue types. Some genes show a marked dispersion of their intermittency, while others have an almost constant probability of being represented by an mRNA transcript. The histogram uses 400 rectangular bins in a 20 × 20 lattice: (a) droplet data, (b) FACS data.

Figure 4 shows a plot of the probability density function (PDF) of ∆*I*_*ik*_ for different tissue types. In each case there is a sharp peak at ∆*I* = 0, indicating a sub-population with very little dispersion. The distribution for the other sub-population has a much greater support, indicating that other genes have probabilities of expression which vary by orders of magnitude between different tissues. The data for the two different experimental protocols are comparable, but not quantitatively equivalent.

**Fig 4.**
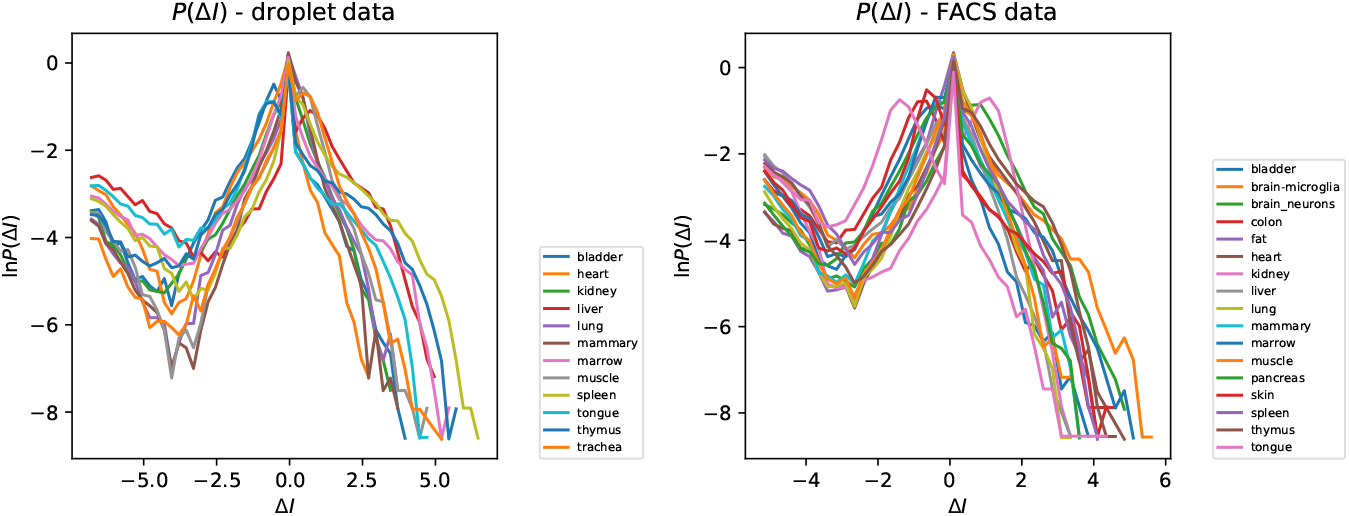
PDF of ∆*I*, representing fluctuations of the intermittency parameter of a gene, for different tissues. There are two populations of genes: the peak at ∆*I* ≈ 0 indicates that some are expressed with nearly the same probability in all tissues, while others vary markedly in their probability of expression: (a) droplet data, (b) FACS data.

The ‘heatmap’ in figure 3 suggests that the peak transcription levels *α*_*i*_ are intrinsic properties of the genes, unaffected by cellular differentiation or by the mechanisms regulating transcription. It might be expected that, at least in some genes, regulation of activity would involve the interaction of transcription factor proteins which could modulate the rate of gene expression by partially blocking the transcription process. If this control molecule were binding and detaching from the DNA on a timescale shorter than the lifetime of mRNA, the gene activity *A* would differ between tissues, which is not observed. However the intermittency of the activity of many genes shows that the timescales for turning transcription on and off are longer than the mRNA lifetime, which is typically taken to be at least one hour, and often considerably longer [12].

### Distribution of gene parameters

We investigated the PDF of the gene activity *A*_*ik*_, illustrated in figure 5, separately for each tissue, showing results for both experimental protocols. Because we have argued that the activity is approximately constant, we might expect that the distribution of the activity *A* will be very similar for different tissues, and the results for the droplet case confirm this. In the case of the FACS data we see curves which are similar, but shifted horizontally, indicating that the overall activity is different in different tissues. Figure 5 also shows the result of applying a ‘normalization’ to make the tissue mean equal to the overall mean. With this tissue-dependent normalization, the distributions of *P* (*A*) from different tissue types are very similar.

**Fig 5.**
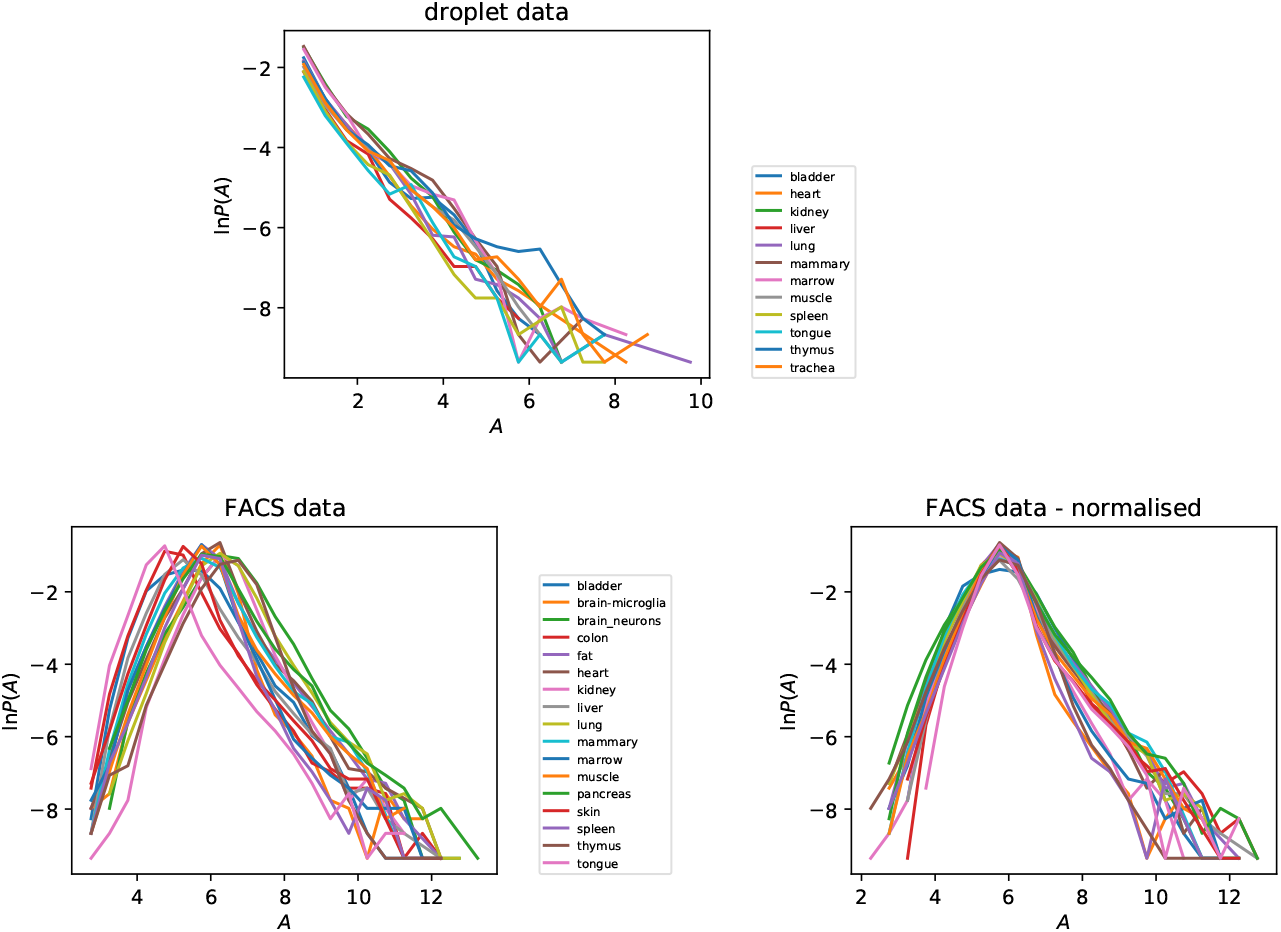
Distribution of gene activity parameter *A*_*ik*_ for different tissues, for both droplet and FACS data. In the case of the FACS data we exhibit the effect of normalizing to equalise the mean activity of different tissues.

In the case of the distribution of the gene activities *A*, there are marked differences between the two different experimental protocols. In particular, in the FACS data there is a ‘tail’ of the distribution corresponding to genes which have very low activities, which is absent from the distribution of activities obtained from the droplet data. The lowest activity genes in the droplet data correspond to events where a single UMI is recorded in a single cell.

We also investigated the PDF of the average of the activity over different genes, 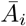 (we shall use an overbar to denote an average over tissue types). The tail of the PDF of 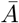, illustrated in figure 6, can be approximated by an exponential function, which corresponds to the PDF of the peak parameter *α* being approximated by a power-law for large *α*: for the droplet data we find *P* (*α*) ~ *α^−^*^2.35^, and *P* (*α*) ~ *α^−^*^2.65^ for the FACS data. The data in figure 6 shows that, while most genes have similar values of the activity parameter, there are a few which have much larger peak activity. We speculate that this can be explained by models in which transcription is impeded by slowly transcribing base sequences. Rare, highly active genes are those which happen not to have the slowly-transcribing sequences.

**Fig 6.**
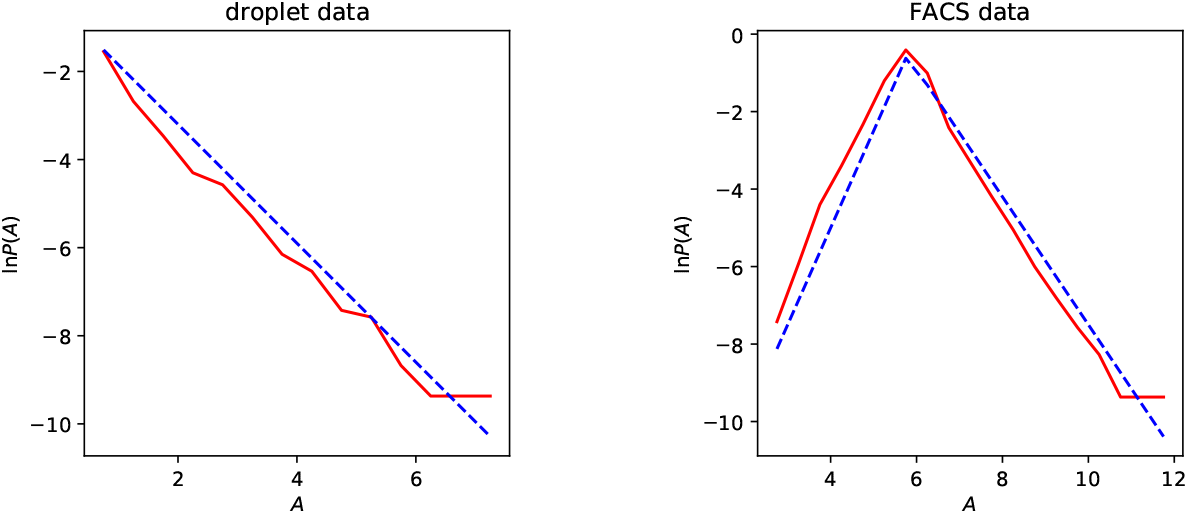
Distribution of tissue-averaged gene activity 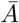 for droplet data and FACS data. The dashed lines are a guide to the eye. The lines in the droplet data plot has slope −1.35, and those in the FACS plot have slopes +2.5 and −1.65.

The PDF of *I* is illustrated in figure 7 separately for each of the tissue types. For each tissue, the PDF of *I* shows a peak close to *I* = 0, corresponding to a sub-population of genes which are expressed with high probability. The distributions of the intermittency *I* do differ between different tissues, in accord with the discussion of section above, but we can say that the PDF of ln(*I*) is very approximately constant over three decades, indicative of a very broad distribution of the intermittency. For the same tissue, the distributions of the intermittency are quite markedly different for the two experimental protocols. This may reflect the greater sensitivity of the FACS experiments.

**Fig 7.**
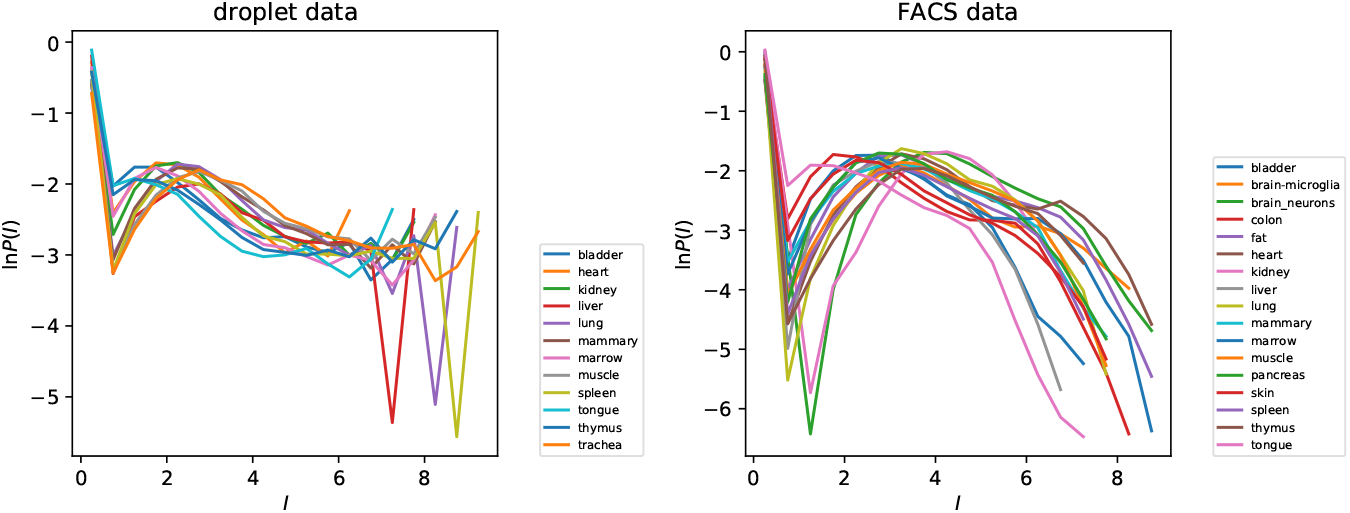
Distributions of gene intermittency parameter *I* by tissue type, for both droplet and FACS data

Figure 7 indicates that a population of genes (with *I ≈* 0) are transcribed continuously, while some others are transcribed with very low probability. The genes which are transcribed with very low probability may be concerned with control functions. The regulatory processes in a cell are probably controlled in a hierarchical manner. The fact that the PDF of ln(*I*) is approximately uniform is interesting, and may be indicative of how the control of transcription is organized.

It might be suspected that genes which have a regulatory function would be required to produce very little protein, and that they might therefore have a low activity parameter *A*, as well as having a high degree of intermittency, *I*. However, the evidence from figure 3 indicates that the expression of genes is regulated via modulation of *I* rather than *A*, indicating that the latter quantity is an intrinsic attribute of the gene, and not susceptible to control. This argument would suggest that there may be no correlation between *I* and *A*. We also examined the correlation coefficient 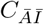 between 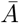 and 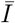, finding 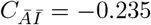 for the droplet dataset and 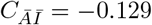 for the FACS dataset, showing that genes with a high intermittency have a slight tendency to have low activity.

## Discussion

Our studies of gene correlations in single-cell mRNA counts for mouse cells distinguish between two possible reasons why some genes are rarely transcribed. They support the view that rarely observed genes are being transcribed intermittently in most cells within a tissue, as opposed to being expressed continuously in specialized sub-populations of cells.

We proposed a characterization of intermittent transcription by assigning two parameters to every gene, namely a peak transcription level *α* and a probability of transcription, *p*. We found that both variables have a very wide range of values, calling for a statistical analysis in terms of logarithmic variables: the activity *A* = ln *α* and the intermittency *I* = − ln *p*.

The wide range of different tissues types in the *Tabula Muris* datasets enable us to gather evidence about the effects of the differentiation of tissues on the transcription process. Single-cell mRNA sequence data provide support for the view that the gene transcription rate *α* in the ‘on’-state is an intrinsic property of the genes, and that differentiation of tissues leads to variation in the probability *p* that transcription occurs. The timescale for turning genes on or off must be slow compared to the lifetime of mRNA molecules (otherwise the level of mRNA would remain nearly constant), but our data do not permit these timescales to be estimated reliably.

This picture of intermittent transcription is consistent with direct observations of ‘bursting’ transcription of genes in other systems. Our results are consistent with the hypothesis that intermittent transcription is ubiquitous in mammalian cells, extending to rarely expressed genes where it would be difficult to observe directly.

Our approach is complementary to studies by Kim and Marioni [10] and Larsson *et al.* [11], who assign parameters to genes based upon a Markovian model for ‘bursting’. We believe that the program of assigning rate coefficients to the bursting model is, however, problematic. Some of the difficulties are

- Single-gene statistics alone cannot distinguish between inhomogeneous counts arising from diversity of cell types and time-dependence of transcription.
- Its interpretation depends upon a particular Markovian, stochastic model of gene transcription which may be over-simplified.
- In some regions of the parameter space, the assignment of kinetic coefficients is highly ill-conditioned. For example, if the rates of binding and unbinding of the control complex are large compared to the rate of degradation of the transcribed mRNA, the amount of mRNA detected depends upon the probability that transcription is occuring, but is insensitive to the timescale of the transcription events.
- In addition to being ill-conditioned, errors arise from the fact that there are additional sources of randomness in the measurement process, resulting from random ‘dropouts’ causing non-detection of some mRNA molecules, and the inherent instability of the polymerase chain-reaction amplification step.

For these reasons, we have presented an alternate approach to making a statistical characterization of transcription, which is more transparent and robust in its interpretation, and which is agnostic about the underlying mechanisms controlling transcription.

The bursting mechanism has been described by stochastic models, based upon telegraph-noise processes [7–9], but these models do not address why stochastic processes appear to play an important role in systems such as mammalian cells where homeostasis is an important principle. Our results suggest two possible reasons why bursting transcription should be ubiquitous.

Intermittent transcription might be used by cells as a matter of necessity, because our results on the constancy of *α* across different tissue types indicate that cells only have ‘on-off’ control, rather than a continuously variable transcription rate. However, beyond being a matter of necessity, intermittent transcription may offer definite advantages over the cellular differentiation model. It makes sense to ‘stage’ operations so that proteins for specific purposes are produced only after their interaction partners are in place. If all the components of complex cellular systems were produced all the time then they would have difficulty finding the complementary sites with which they should bind. Limits on the rate of ribosomal translation imply that a cell cannot be efficiently performing all of its functions all of the time.

These arguments about using intermittent transcription to organize the efficient operation of a cell would suggest that, rather than modeling bursting as a purely stochastic phenomenon, we should look for a model of bursting transcription which is determined by a complex dynamical system which can turn genes on and off in a time-ordered sequence.

Finally, we remark that studies of post-transcriptional modification of mRNA should be capable of yielding additional information about the timescales of the intermittent gene transcription, by an extension of the approach described in [17].

## Acknowledgments

We thank Joshua Batson, Olga Botvinnik and Emma Lundberg for comments. MW thanks the Chan Zuckerberg Biohub for its hospitality. GH thanks the Kavli Institute for Theoretical Physics (supported by grant NSF-PHY-1748958) for its hospitality.

## Notes

#### Summary of Updates

Minor revisions and corrections in response to comments, no substantive changes.

